# Unveiling Tryptophan Dynamics and Functions Across Model Organisms via Quantitative Imaging

**DOI:** 10.1101/2024.02.12.580012

**Authors:** Kui Wang, Tian-lun Chen, Xin-xin Zhang, Jian-bin Cao, Pengcheng Wang, Mingcang Wang, Jiu-lin Du, Yu Mu, Rongkun Tao

**Affiliations:** Institute of Neuroscience, State Key Laboratory of Neuroscience, Center for Excellence in Brain Science and Intelligence Technology, Chinese Academy of Sciences, 320 Yue-Yang Road, Shanghai 200031, China; University of Chinese Academy of Sciences, 19A Yu-Quan Road, Beijing 100049, China; School of Life Science and Technology, ShanghaiTech University, 319 Yue-Yang Road, Shanghai 200031, China; Department of Anesthesiology, Taizhou Hospital of Zhejiang Province affiliated to Wenzhou Medical University, 150 Xi-men Road, Zhejiang 317000, China; Department of Pediatric Cardiology, Xinhua Hospital, Shanghai Jiao Tong University School of Medicine, 1665 Kong-Jiang Road, Shanghai 200092, China

**Author notes:** These authors contributed equally to this work (Kui Wang, Tian-lun Chen, Xin-xin Zhang, Jian-bin Cao).

## Abstract

Tryptophan is an essential amino acid involved in many cellular processes in vertebrates. Systemic and quantitative measurement of tryptophan is crucial for evaluating its essential role as a precursor of serotonin and kynurenine, key neuromodulators affecting neural and immune functions. We utilized a robust and highly responsive ratiometric indicator for tryptophan (GRIT) to quantitatively measure tryptophan dynamics in bacteria, mitochondria of mammalian cell cultures, and human serum. At the cellular scale, these analyses uncovered differences in tryptophan dynamics across cell types and organelles. At the whole-organism scale, we revealed that inflammation-induced tryptophan concentration increases in zebrafish brain led to elevated tryptophan metabolites serotonin and kynurenine levels, which is associated with the prolonged sleep duration. The reduction of zebrafish plasma tryptophan was mirrored in patients with inflammation symptoms and could serve as a biochemical marker of inflammation. In summary, this study introduces GRIT as a powerful method for studying tryptophan metabolism and functions across scales and species and suggests that tryptophan metabolic processes link the immune response and animal behavior.

## INTRODUCTION

Tryptophan, an essential amino acid exclusively obtained from food, plays a crucial role in cellular protein synthesis and survival. Its absorption, transportation, and metabolic pathways within all vertebrates are highly conserved, leading to the production of various active molecules through pathways involving kynurenine, serotonin, and indole^1-3^. Thus, tryptophan metabolism serves as a pivotal metabolic regulator for processes such as immune responses, neural activities, and intestinal functions.

The tryptophan level metabolism is regulated at subcellular level, cell types, and across systems^4^. The abnormalities in tryptophan metabolism can induce systemic effects, influencing other tissues via the circulatory system, potentially triggering or exacerbating diseases like neurodegeneration, infections and cancer^5^. Moreover, tryptophan levels in plasma are closely associated with disease progression and prognosis in clinic^6^. Such extensive implications highlight the importance of systematically quantifying tryptophan dynamics across organisms.

Traditional methods, such as high-performance liquid chromatography (HPLC), are limited in revealing metabolic differences between different cell types and among subcellular compartments, also lack the precise temporal resolution to track rapid cellular metabolic changes during cellular stress^7^. In contrast, our recently developed genetically encoded tryptophan sensor, GRIT^8^, represents a significant advancement over FRET-based tryptophan indicators (FLIPW-CTYT)^9^. GRIT offers a substantially larger dynamic range (9.55 versus 0.3), a smaller molecular size (39.7 kDa versus 80.1 kDa), and reduced photobleaching. These improvements make GRIT a highly effective tool for the quantitative measurement of tryptophan.

In this study, we demonstrate GRIT allows a new paradigm offering high spatiotemporal resolution, systematically quantifying dynamic changes in quantitatively mapping tryptophan dynamics in bacteria, mitochondria, zebrafish and human serum. Such advancements can facilitate a comprehensive understanding of the differences and dynamics in metabolite levels among cells within different tissues under physiological and pathological conditions, and reveals tryptophan metabolic processes link the immune response and animal behavioral changes.

## RESULTS

### Monitoring tryptophan dynamics in living bacteria

Bacteria possess a tight regulatory mechanism on tryptophan metabolism^10^. We investigated tryptophan uptake into *Escherichia coli* cells by expressing the GRIT sensor in the cytoplasm (**Figures 1A and 1B**). Exogenous tryptophan addition induced a strong and rapid fluorescence response in GRIT-positive bacteria, with a maximum ∼3-fold increase (**Figures 1B and 1C**), but not in GRITOL-expressing bacteria (**Figure 1B**). GRIT could respond to 0.1 - 100 μM exogenous tryptophan in a dose-dependent manner with an absorption rate of ∼16.5 μM/s (**Table S1**). The Michaelis constant (*K*_m_) of tryptophan uptake (∼0.58 μM) (**Figure 1D**) is similar to the reported biophysical properties of aroP (∼0.4 μM)^11^, a tryptophan transporter that is highly expressed in bacteria according to the transcriptome^12^ (**Figure S1A**). The free concentrations of intracellular tryptophan in tryptophan deprivation and tryptophan exposure conditions were measured to be 10.6 ± 3.7 μM (similar to the reported level 12 μM^13^) and 192 ± 23.9 μM, respectively (**Table S1**). In addition, no significant fluorescence change was observed when 19 other tested amino acids were added in the absence or presence of tryptophan (**Figure S1B**). These results suggest that bacterial tryptophan uptake is highly sensitive and specific.

**Figure 1.**
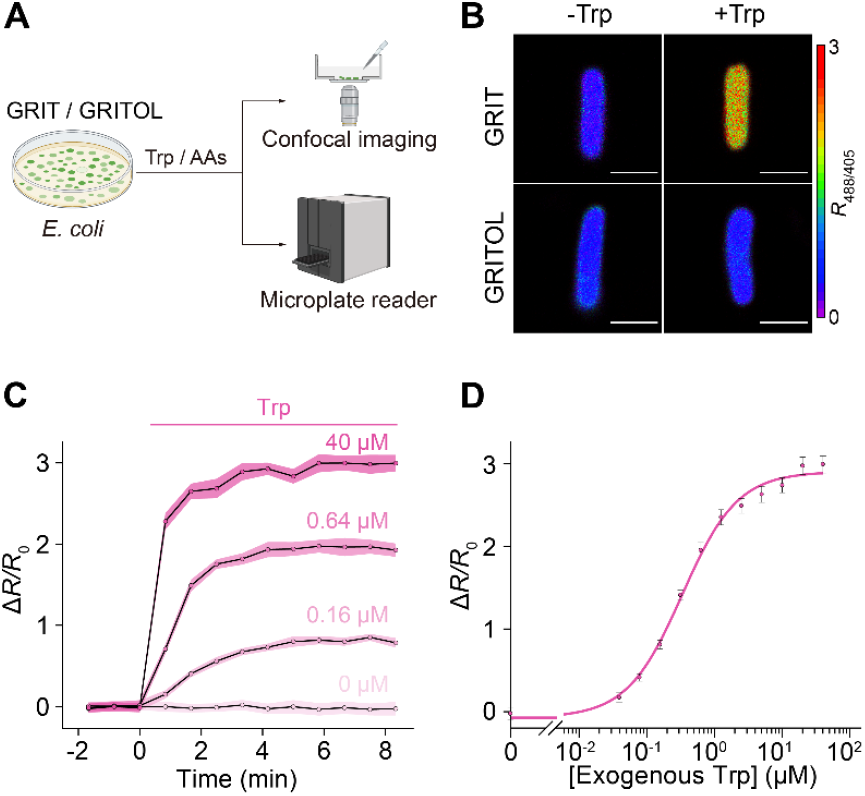
Detection of tryptophan uptake in bacteria with GRIT sensor. **(A)** Schematic representation illustrating the fluorescence detection of tryptophan (Trp) dynamics with the GRIT or GRITOL sensor in *Escherichia coli* cells. **(B)** Fluorescence images of bacteria expressing GRIT and GRITOL upon the addition of 0.1 mM Trp. **(C)** Changes in the fluorescence of GRIT in response to various concentrations of Trp in *Escherichia coli*. **(D)** The dose-response curve of GRIT-positive bacteria treated with exogenous Trp. Data are from panel Fig. 1C. Scale bars, 1 μm. Data shown as mean ± s.e.m., n = 3 independent experiments in C and D. See also **Figure S1**, and **Table S1**.

### Detection of mitochondrial tryptophan dynamics in mammalian cells

Mitochondria are semiautonomous subcellular organelles with their own genome and require tryptophan transport from the cytosol to synthesize mitochondrial proteins. However, due to the lack of specific detection methods, reports on mitochondrial tryptophan metabolism have been sparse^14^. We selectively targeted the GRIT sensor in the mitochondrial matrix (**Figures 2A and 2B**), resulting in a marked increase in fluorescence signals compared to the control cells (**Figures S2A and S2B**). This enhancement enables effective fluorescence measurement using both microplate readers and microscopy (**Figure 2A**). Mito-GRIT showed an ∼1.7-fold fluorescence increases upon the addition of 0.5 mM tryptophan (**Figure 2B**). The mitochondrial tryptophan concentration was determined to be 112.4 ± 10.0 μM **(Table S1)**, which was close to the HPLC measurements (108.0 ± 10.3 μM, **Table S1**). However, the tryptophan uptake rate of mitochondria (∼9.6 μM/min) was only ∼18% of that in the cytoplasm (∼52.6 μM/min) (**Figure 2C, Table S1**). Upon tryptophan deprivation, the mitochondrial tryptophan level was relatively stable in the first 20 min and decreased linearly at a rate of ∼0.8 μM/min (**Figure 2C, Table S2**). This suggests that the mitochondrial tryptophan pool is predominantly regulated by cytosolic supply and the mitochondrial consumption, such as protein synthesis.

**Figure 2.**
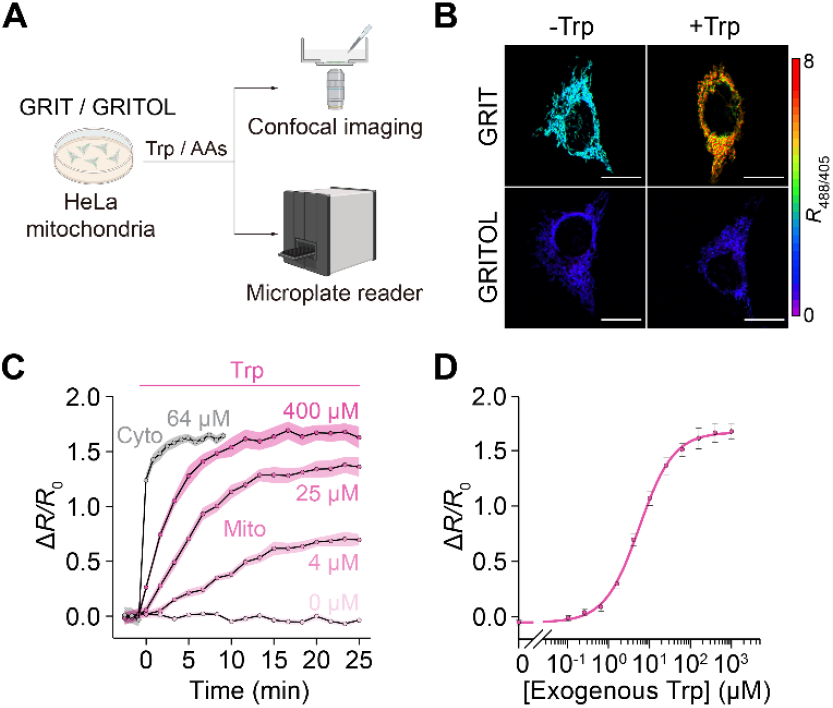
GRIT sensor reports mitochondrial tryptophan dynamics in cultured HeLa cells. **(A)** Schematic showing the fluorescence detection with mito-GRIT or mito-GRITOL sensor in HeLa cells. **(B)** Fluorescence responses of HeLa cells expressing GRIT (upper) and GRITOL (bottom) in mitochondria upon the addition of exogenous 0.5 mM Trp in HBSS buffer. **(C** and **D)** The traces (C) and dose-dependence curve (D) of mitochondrial GRIT sensor in response to varying concentrations of added tryptophan. The gray trace in C represents the fluorescence response kinetics of cytosolic GRIT to indicated Trp concentration (64 μM) in HBSS buffer, suggesting a significantly faster tryptophan uptake into the cytosol compared to the mitochondria. Scale bars, 10 μm. Data shown as mean ± s.e.m. n = 3 independent experiments. See also **Figure S2** and **Tables S1, S2**.

We have demonstrated that cytosolic tryptophan pool is finely regulated by the supply of external amino acids, which can be classified into three groups, Class I (large and neutral amino acids, classic LAT1 substrate), Class II (five uncharged amino acids), Class III (six charged or nonpolar amino acids)^8^. To further understand the characteristics of mitochondrial tryptophan transportation, we detected mitochondrial tryptophan dynamics in response to all other 19 amino acids. The six amino acids from Class III, which have no effect on cytosolic tryptophan flux, also did not affect mitochondrial tryptophan dynamics (**Figures S2C-E and Table S2**). In contrast, the 13 amino acids from Class I and Class II, despite their varied effects on cytosolic tryptophan efflux, displayed similar impacts on mitochondrial tryptophan dynamics, both in the absence or presence of tryptophan (**Figures S2C-E and Table S2**). Collectively, the biophysical properties of mitochondrial tryptophan metabolism suggest specific and unidirectional mitochondrial tryptophan transporters, which are different from the bidirectional transportation of cell surface LAT1^8^.

### Detection of serum tryptophan in patients with inflammation syndrome

The clinical relevance of tryptophan metabolism lies in its association with various diseases and inflammatory responses, particularly evident in the variations in serum levels^6^. Therefore, a rapid and efficient *in vitro* detection method could significantly advance diagnostic technologies. We demonstrated that purified GRIT sensor could show large fluorescence ratio changes to tryptophan ranging from 20 μM to 1 mM (**Figure S3A**). This broad detection range is line with the theoretical prediction of an established web application “SensorOverlord”^15^ (**Figure S3B**), which overlaps with the concentration fluctuations of tryptophan in most physiological and pathological conditions.

We collected serum samples from patients with inflammatory symptoms characterized by following specific clinical criteria^16^: hypersensitive C-reactive protein (hs-CRP) concentration > 50 mg/L, and neutrophil ratio > 80%, white cell count > 10,000/μL (**Figure 3A**). We first analyzed serum tryptophan levels using HPLC assay, followed by GRIT sensor detection of the remaining portion of the same sample (**Figure 3A**). Despite the autofluorescence of blood may interfere with the fluorescence assays, our results showed that GRIT sensor possess much higher fluorescence signal than the serum background (**Figures S3C-F**). In addition, the quantified tryptophan concentration was highly consistent between HPLC and GRIT sensor measurements (**Figure 3B**), demonstrating the robustness of tryptophan measurement with purified GRIT sensor in human serum samples. The serum tryptophan level in inflamed patients (44.7 ± 4.7 μM) decreased ∼25% compared with that of control (59.6 ± 3.6 μM) (**Table S1**). This reduction in serum tryptophan was consistently observed under various conditions, including different sexes, ages, and extents of inflammation (**Figures 3C-3F, S3G-I**). These results validate the GRIT sensor as an alternative high-throughput *in vitro* diagnostic method and a decrease in serum tryptophan level could be a biochemical marker of inflammation in patients.

**Figure 3.**
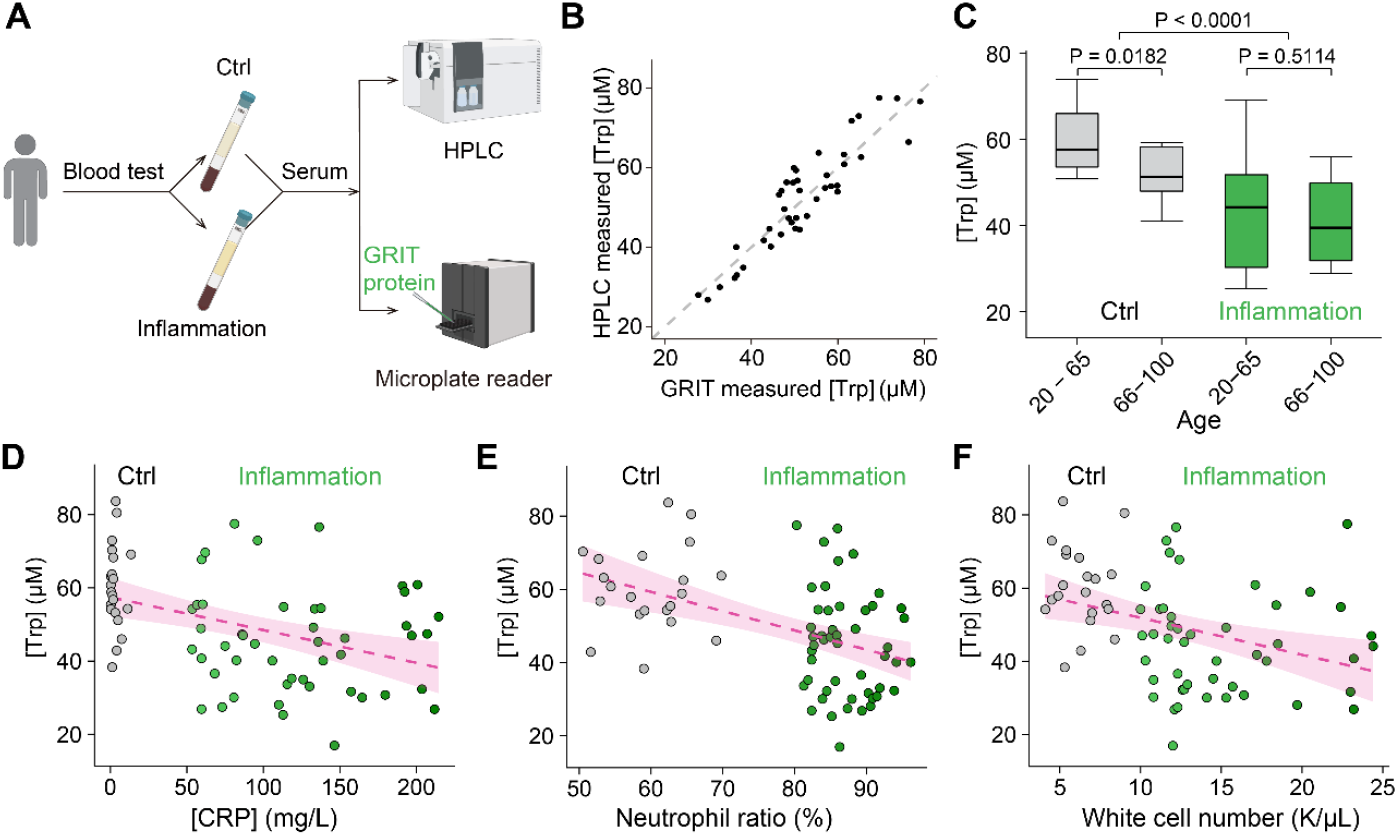
Decreased serum tryptophan concentration is a biochemical mark of inflamed patients. **(A)** Schematic of the experimental procedures for the quantification of human serum tryptophan. **(B)** A comparative analysis of quantitative serum tryptophan level in identical samples measured by GRIT sensor assay or HPLC measurement (n = 42). **(C)** Tryptophan concentrations of human serum samples in the control group (grey, n = 12 in 20-65 subgroup, n = 13 in the 66-100 subgroup) and inflammation group (green, n = 17 in 20-65 subgroup, n = 35 in the 66-100 subgroup) based on age. Note that the total n number (42) in panel B are smaller than the n number (77) in C, due to the volume of some serum samples were not enough for GRIT sensor assay after HPLC (< 0.3 mL). (**D-F**) Relationships between tryptophan concentrations in human serum samples and concentrations of hypersensitive C-reactive protein (D), neutrophil ratio (E) and white cell number (F). The dashed magenta line represents a linear fit of the data, with the shaded area indicating the fitting confidence interval. Two-tailed student’s unpaired t test for C. See also **Figure S3** and **Table S1**.

### Detection of tryptophan metabolites and behaviors during inflammation in animal model

In the central nervous system, tryptophan is catabolized into serotonin and kynurenine, both of which modulate neural activities and are related to sleep, depressive behaviors, and fatigue symptoms^17-20^. To further explore the pathological implications of the documented inflammation-triggered tryptophan redistribution phenomenon^8^, we subsequently exposed zebrafish to lipopolysaccharide (LPS). This resulted in a dose-dependent increase in sleep duration (**Figures 4A-C and S4A**), consistent with the sustained elevation of tryptophan levels in the brain following LPS treatment (**Figures S4B and S4C**). Next, we measured the concentrations of tryptophan, serotonin, and kynurenine in isolated zebrafish brains. Inflamed zebrafish showed a slight but non-significant increase in total tryptophan, a significant over 3-fold increase in serotonin, and a 36% increase in the kynurenine, compared to control individuals (**Figures 4D and 4E**). The rises of tryptophan metabolites in the brain may contribute to the increased sleep behaviors during inflammation. Furthermore, inhibiting these metabolic pathways resulted in developmental deformities, leading to their demise (**Figures S4D and S4E**), suggesting that disrupting tryptophan metabolism leads to severe pathological consequences.

**Figure 4.**
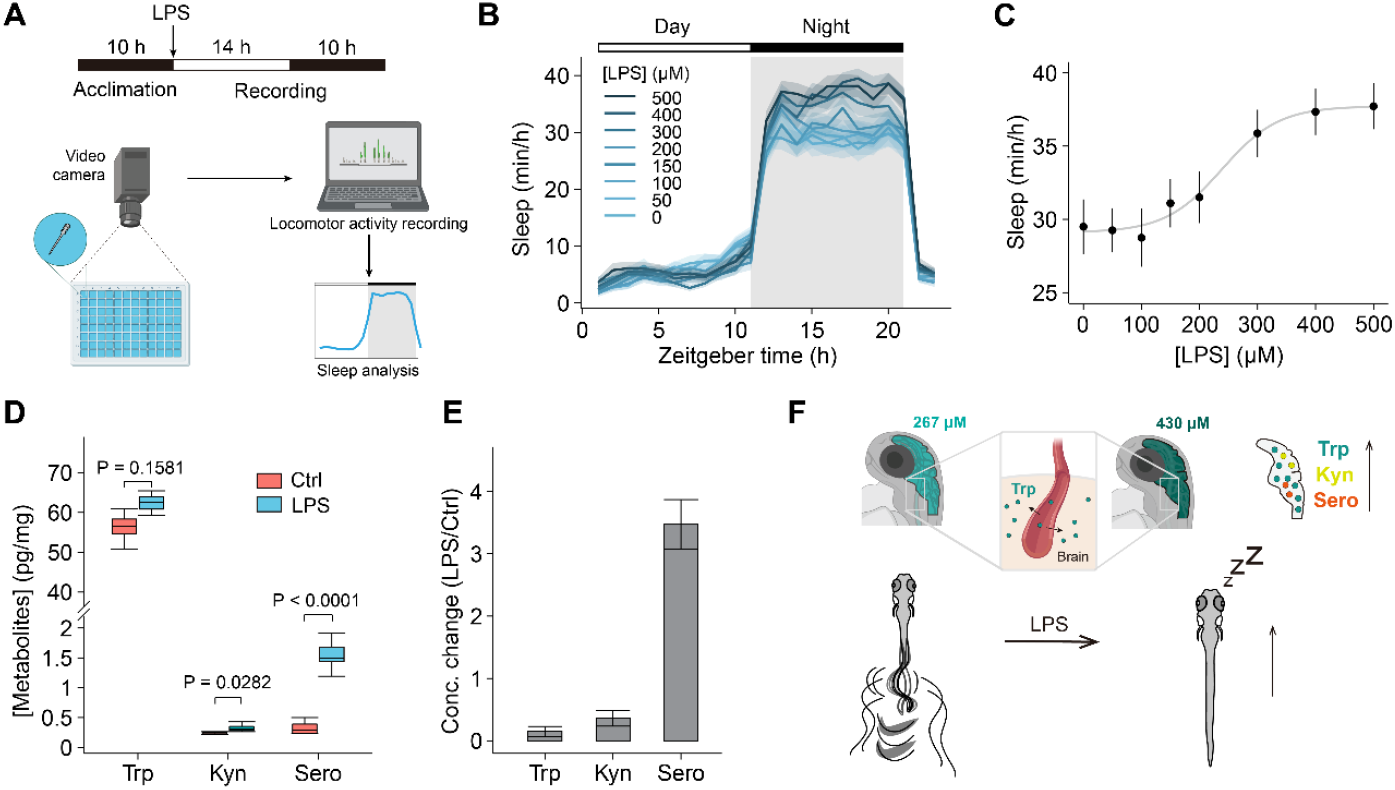
Detection of metabolites levels in zebrafish larvae and zebrafish sleep changes during inflammation. **(A)** Schematic of the zebrafish sleep behavior measurement during inflammation. **(B)** Changes in the sleep time of zebrafish in response to various concentrations of LPS (n = 60 fishes from 5 - 7 independent experiments). LPS was added at 0 hr with indicated concentrations. **(C)** The dose-response curve of mean sleep time at night of zebrafish treated with different concentrations of LPS. **(D-E)** The absolute levels (pg/mg) (D) and relative changes (E) of tryptophan, kynurenine and serotonin in isolated zebrafish brain with (n = 7) or without LPS treatment (n = 6). **(F)** The proposed working model of the dynamics and functions of tryptophan in zebrafish during inflammation. During inflammation, plasma tryptophan enters the brain, elevates the serotonin and kynurenine levels, and increases zebrafish sleep time. The concentrations of brain tryptophan concentrations were adapted from our previous study^8^. Data shown as mean ± s.e.m. Two-tailed unpaired student’s t test for D. See also **Figures S4**.

In summary, our results demonstrate that inflammatory responses modulate tryptophan levels across tissues (**Figure 4F**), with altered metabolic levels in the brain influencing behavior. This indicates a metabolic link between the inflammatory response and animal behavior.

## DISCUSSION

In this study, we have demonstrated the versatility of using GRIT biosensor for quantitatively measuring tryptophan levels across divers biological models, including bacteria, subcellular organelles, and human serum. The GRIT, a superior genetically encoded tool, supports multiple applications and long-term recordings of tryptophan.

Our findings show that intracellular tryptophan levels of different cell types and animal models typically lie in the sub-millimolar range with distinct transportation properties (**Supplementary Table 1**), which are in relevance to their specific functions in respective cells. In *Escherichia coli*, tryptophan anabolism stands out as one of the most energy-intensive metabolic pathways^21^, which is activated exclusively under tryptophan deprivation conditions (lower than sub-micromolar), along with a significant increase in its half-proliferation time^22^. The high affinity and rapid response to tryptophan facilitate rapid growth to adapt variable environments^22^ and allow gut microbes to continually metabolize tryptophan into indole derivatives, influencing host the enteric nervous systems^23,24^.

In mammal system, reprogramming tryptophan metabolism has become attractive for tumor therapy, such as targeting LAT1 or indoleamine 2,3-dioxygenase (IDO)^5,25^. In contrast to the bidirectional and rapid turnover of the cytosolic tryptophan pool, mitochondrial tryptophan transportation exhibits a slower rate and higher specificity. The stringent regulation of mitochondrial tryptophan pool may benefit protein synthesis and quality control within mitochondria. Considering that the intense, ratiometric GRIT sensor is suitable for high-throughput screening, it offers an opportunity to identify the mitochondrial transporter^26^, and an optical method and paradigm for studying the metabolism of subcellular organelles.

Quantitative measurements by GRIT sensor in intact animal and human serum could reveal systemic tryptophan dynamics and its functions. Tryptophan metabolism regulates immune systems and neural behaviors through kynurenine and serotonin pathways^5,19^,27, making the redistribution of tryptophan crucial for balancing the physiological functions under pathological conditions^4,28^. Our measurements suggest that a flow of plasma tryptophan to the brain and the consequent increase of tryptophan metabolites (serotonin and kynurenine) during inflammation, resulted in prolonged sleep time in zebrafish. These results align with the elevated serotonin concentration in the brain and fatigue and depression behaviors in LPS-treated mice^29^ and patients with inflammation symptoms, suggesting a potential metabolic link between immune response and animal behavior^1,48^.

*In vitro diagnostics* plays a pivotal role in disease detection, monitoring, and personalized medicine, ultimately contributing to better patient outcomes and overall public health^30,31^. The simple, fast and accurate GRIT probe allows high throughput quantification of tryptophan concentrations in a patient’s serum, which marks a significant advancement in the clinical applications of fluorescent probes. The reductions in serum tryptophan levels during inflammatory responses exhibit remarkable consistency between zebrafish and human patients. Interestingly, a similar redistribution of tryptophan has been observed in glioblastoma^32^. Moreover, our results suggest a correlation between patients’ ages and serum tryptophan levels (**Figures 3C, S3H and S3I**), but not with gender, implying a potential role of tryptophan in aging, which are consistent with a previous report that tryptophan supplement could extend worm lifespan via mitochondria NAD *de novo* synthesis^8^. These results underscore the highly conserved nature of tryptophan metabolism and circulation among vertebrates, emphasizing the relevance of our observations across different species. Considering inflammation is a hallmark of various diseases, such as cancer, aging, obesity, and cardiovascular disease, the systemic and quantitative measurement of tryptophan dynamics on zebrafish offers new insights into the metabolic regulation mechanisms of these diseases^4,5,33.^

Overall, the systemic quantification of tryptophan metabolism with the GRIT sensor portends the direct imaging of intracellular tryptophan metabolism and further understanding of its functions under both physiological and pathological conditions. Quantitative imaging paradigms in model animals is expected to pave the way for new therapeutic approaches in human metabolic diseases.

## STAR METHODS

Detailed methods of this paper include the following:

KEY RESOURCES TABLE

RESOURCE AVAILABILITY

Lead contact

Data availability

EXPERIMENTAL MODEL AND SUBJECT DETAILS

Cell cultures

Zebrafish

Human plasma samples

METHOD DETAILS

Plasmids construction

Cell culture, transfections, and establishment of stable cell lines

Fluorescence measurement of living cells with microplate reader

Fluorescence microscopy

*In vitro* transcription and mRNA purification

Zebrafish Sleep and Locomotion Analysis

Human plasma sample preparation

Quantitative measurement of tryptophan with GRIT sensor *in vitro* and *in vivo*

Tryptophan quantification with high performance liquid chromatography

Statistical Analysis

## ACKNOWLEDGMENTS

We thank Dongjian Wang, Hongbo Chi, Haifei Xiang, Zhehang Zhou,Xiandong Shao, Guo Yu, Rui Lu, Yuqian Cai for their kind supports and suggestions of patient inflammation test. We appreciate the technical assistance from Dr. Hanyang Hu, Dr. Xufei Du, Hao Deng and Dr. Weixi Feng. We thank Jiwen Bu for secretarial assistance. We thank Chemical Biology Core Facility and Molecular Biology Core Facility in CEMCS, CAS for technical support. This work was supported by grants from National Science and Technology Innovation 2030 Major Program (2021ZD0202203, SQ2021AAA010882,2021ZD0204500, 2021ZD0203704), National Natural Science Foundation of China (21171090, 32171026, 32321003), Shanghai Science and Technology Commission (21ZR1482600), Shanghai Municipal Science and Technology Major Project (2018SHZDZX05), 2023 Youth Innovation Promotion Association CAS, Strategic Priority Research Program of the Chinese Academy of Sciences (XDA27010403), Scientific Instrument Developing Project of the Chinese Academy of Sciences (YJKYYQ20210029), and Basic Public Welfare Research Project of Zhejiang Province (LGD20H090005).

## AUTHOR CONTRIBUTIONS

R.T. and Y. M. conceived the project. R.T., K. W., T. C., J. C. and X. Z. performed experiments. R.T., Y. M., K. W., T. C. and X. Z. analyzed data and made figures. R.T., Y. M. and J. D. wrote the draft and revised the manuscript.

## DECLARATION OF INTERESTS

The authors declare no competing financial interests.

## STAR METHODS

### KEY RESOURECES TABLE

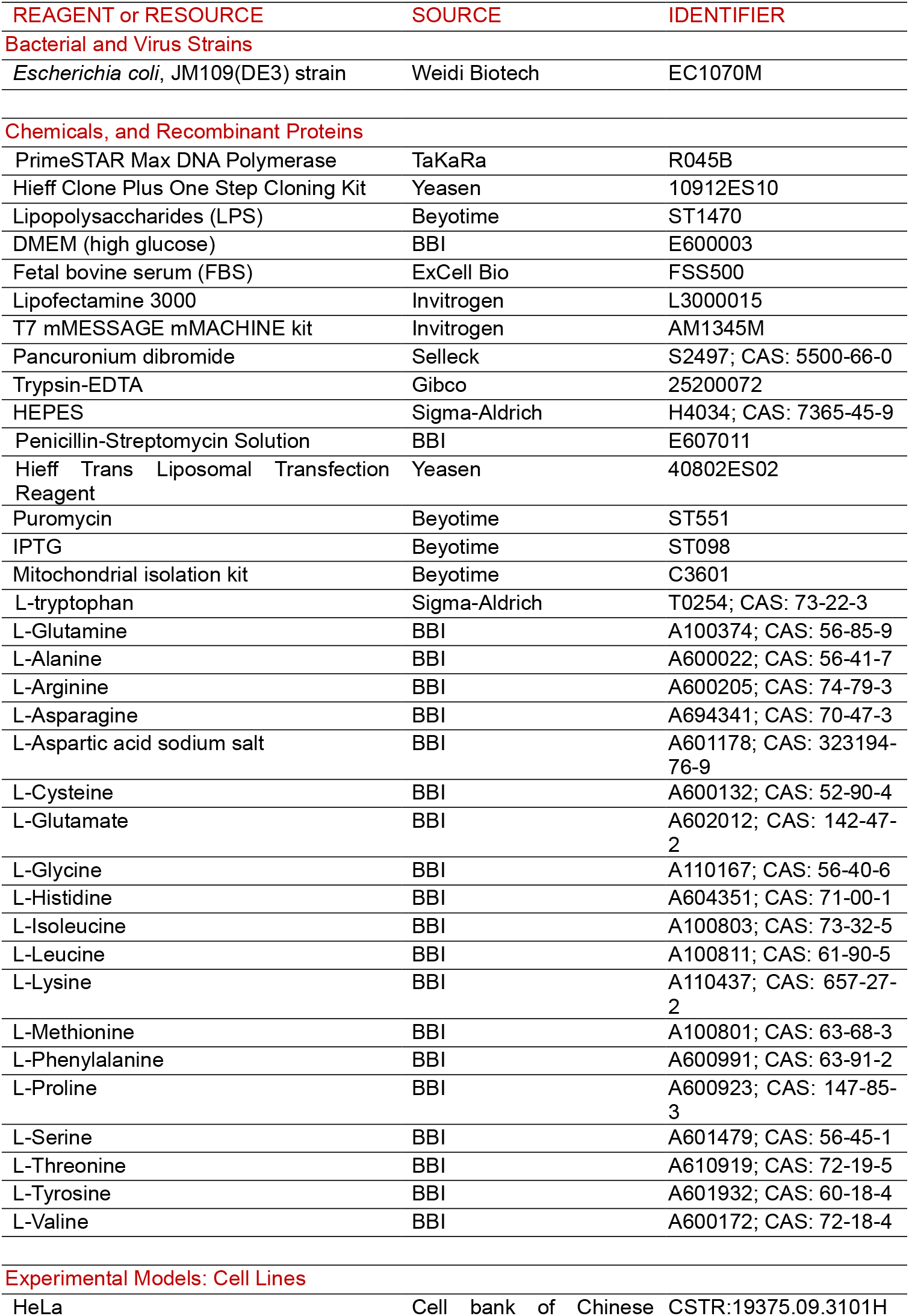
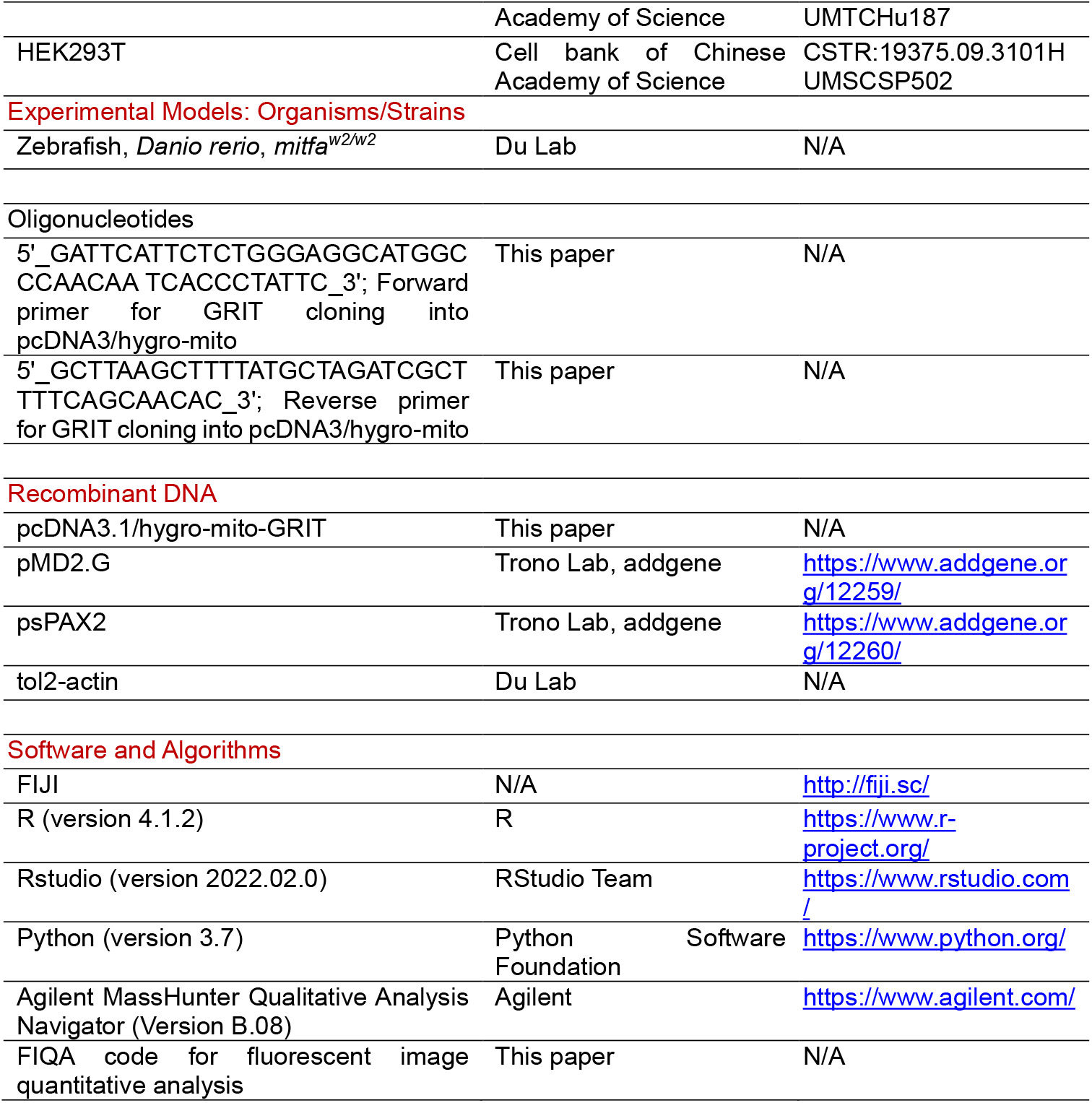

### RESOURCE AVAILABILITY

#### Lead contact

Further information and requests for resources and reagents should be directed to and will be fulfilled by the Lead Contact, Rongkun Tao.

#### Data availability

The data that supports the conclusions of this study are available from the corresponding author upon request. GRIT and GRITOL constructs for mammalian expression are available on the WeKwikGene plasmid repository at Westlake Laboratory, China (https://wekwikgene.wllsb.edu.cn/). All other constructs are available upon requests.

### EXPERIMENTAL MODEL AND SUBJECT DETAILS

#### Cell cultures

HeLa and HEK293T cell lines, obtained from the Cell Bank of the Chinese Academy of Science, were confirmed to be free of mycoplasma contamination. These cell lines were cultured in high glucose DMEM (BBI), supplemented with 10% FBS (ExCell Bio) and 1% penicillin-streptomycin (10,000 U/ml). The cells were maintained at 37 °C in a humidified atmosphere composed of 95% air and 5% CO_2_.

#### Zebrafish

Colonies of Zebrafish (*Danio rerio*), specifically the Nacre strain (*mitfa*^*w2/w2*^), were maintained following standard procedures. The experimental protocols involving zebrafish were approved by the Chinese Academy of Sciences (Approval no. NA-046-2019). The zebrafish were kept under a light-dark cycle of 14 hours light to 10 hours dark, and the water temperature was consistently maintained at 28 °C.

#### Human plasma samples

All the population studied were from patients of Taizhou hospital of Zhejiang Province affiliated to Wenzhou Medical University. The experiments were performed according to the medical ethics rules (no. K20211202).

## METHOD DETAILS

### Plasmids Construction

For mitochondrial expression, the mitochondrial signal peptide from CoxVIII (cytochrome c oxidase subunit VIII) was duplicated and fused at the N-terminus of tryptophan biosensors^7^. To make lentiviral vectors, genes of mito-GRIT and mito-GRITOL were subcloned into pLVX-IRES-Puro backbone by NheI and PmeI.

### Cell Culture, Transfections, and Establishment of Stable Cell Lines

To generate lentiviral particles, HEK293T cells were transfected with lentiviral plasmids, including two packaging vectors (pMD2.G and psPAX2), using Hieff Trans (YEASEN). Lentivirus-containing supernatants were collected at 48 and 72 hours post transfection and subsequently applied to HeLa cells cultured in 6-well tissue culture plates supplemented with 8 μg/ml polybrene. Stable HeLa cell lines were established by cultivating the cells with 1 μg/ml puromycin for one week, followed by sorting using MoFlo Astrios EQ (Beckman Coulter) with a 488 nm laser line. The resulting stable HeLa cells were used for subsequent fluorescence measurement experiments.

### Fluorescence Measurement of Living Cells with Microplate Reader

Fluorescent bacteria cells were centrifugated at 4000 rpm for 2 min, followed by washing three times with modified PBS buffer containing 10 mM HEPES, pH 7.3. The optical density (OD_600_) of bacterial suspensions was diluted to ∼0.01, followed by a 30 min pre-starvation before measurements. Three points were detected as the steady state, followed by addition of different concentrations of tryptophan and a 10-min continuous reading on the BioTek SynergyNeo Multi-Mode microplate reader. The fluorescence ratios (*R*_485/420_) of GRIT and GRITOL were divided to fit with titration curve for calibration.

The HeLa cell lines stably expressing GRIT sensors were trypsinized, resuspended and plated in a black 96-well flat-bottom plate at a density of 26,000 cells per well, allowing them to cultivate for 10-12 hours. Before the fluorescence measurements, the DMEM media was removed, and cells were washed twice with freshly prepared HEPES buffer. Various amino acids were then introduced into the detection buffer, each at specified absolute concentrations. To correct the fluorescence values of GRIT sensors, the signals from HeLa cell samples were background-subtracted.

### Fluorescence Microscopy

Fluorescent bacteria cells were collected and diluted to the OD_600_ ∼0.2 with modified PBS buffer with or without 0.1 mM tryptophan. Bacteria solution (10 μL) was added on the 0.17 mm coverslips (Nunc), then the coverslip was put upside down on a glass slide for imaging. Cultured HeLa cells expressing Mito-GRIT and Mito-GRITOL were transferred to an 8-well glass-bottomed dish (Cellvis) where the culture medium was replaced with HEPES buffer with or without 0.5 mM tryptophan prior to experimentation. Utilizing an inverted confocal Olympus FV3000 automatic microscope with a UPlanSApo 40 × Sil objective (N.A. 1.25), dual-excitation ratio imaging was performed. The GRIT sensor was excited using 405 nm and 488 nm lasers, capturing emission figures between 500-550 nm with a photomultiplier tube (PMT) in a 1024×1024 format and 12-bit depth. For subsequent data analysis, a semiautomatic analysis was carried out using the FIQA code implemented in ImageJ Fiji. This analysis involved several crucial steps, including background subtraction to eliminate unwanted noise, skin removal for enhanced clarity, generation of ratiometric pseudocolored images to visualize variations, and batch measurement of fluorescence intensities.

### *In Vitro* Transcription and mRNA Purification

The mRNA preparation involved the utilization of a T7 mMESSAGE mMACHINE kit (Invitrogen). We amplified the cDNAs corresponding to GRIT, GRITOL, or Tol2 and performed *in vitro* transcription following the manufacturer’s guidelines. To obtain purified mRNA, a two-hour incubation in lithium chloride at -20 °C was performed, followed by dilution with DEPC-treated nuclease-free water. This approach facilitated the generation of high-quality mRNA samples for further experimental applications.

### Zebrafish Sleep and Locomotion Analysis

Sleep analysis was performed based on tracking results of individual larval zebrafish. Zebrafish larvae were raised with a light (14 hrs) and dark (10 hrs) cycle. The daytime starts at 9 am and the night-time starts at 11 pm. When experiments started, individual larvae were transferred into wells of a 96-well plate, which was placed inside Zebrabox (Viewpoint Life Sciences, Lyon, France). Their locomotor activity was recorded using the quantization mode in a video-tracking system (Viewpoint Life Sciences). Parameters for detecting different movement patterns were determined empirically: detection thresholds, 15; burst, 29; freeze, 3; bin size, 60 s. PBS or different doses of LPS were added into the bath at 9 am on the second morning and maintained throughout the experiment.

### Human Plasma Sample Preparation

Venous blood samples of patients were collected in tubes early in the morning, after an overnight fast for a standard blood test, which are divided into control group and inflammation group based on the blood test with clinical criteria^16^. For control group: (1) the number of white cells is between 4,000 – 9, 000/μL; (2) the concentration of hypersensitive C-reactive protein (hs-CRP) is less than 20 mg/L; (3) the neutrophil ratio is between 50% - 70%. For inflammation group: (1) the number of white cell is over 10, 000/μL; (2) the concentration of hs-CRP is more than 50 mg/L; (3) the neutrophil ratio is over 80%. The studied population consisted of 77 adult subjects, which were randomly selected independent of gender and age. Selected plasma samples were centrifuged at 3,500 rpm for 2 min and kept at 4 °C until analysis.

### Quantitative Measurement of Tryptophan with GRIT sensor *In Vitro* and *In Vivo*

Accurate and quantitative assessments of tryptophan levels in cells and serum samples are achievable through the calculation of normalized sensor fluorescence ratios and comparing these ratios with those obtained from purified GRIT protein in vitro, as described previously^8^.

For the quantification of serum tryptophan concentrations with GRIT sensor, serum sample was equally mixed with 1 μM GRIT or GRITOL sensor in 100 mM HEPES buffer, 100 mM KCl (pH 7.3), followed by routine fluorescence measurement similar to protein titration experiments.

For quantification bacterial tryptophan, we measured the excitation fluorescence ratios of cells expressing the GRIT sensor with or without 40 μM tryptophan as the intracellular tryptophan levels in tryptophan-fed cells and tryptophan-starved cells, respectively. To account for the pH effect, the excitation ratios of cells expressing GRIT were corrected by those of cells expressing GRITOL measured in parallel and fitted with an *in vitro* protein titration curve at pH 7.3.

For *in situ* live-cell calibration, the unbound and saturated states of the mito-GRIT sensor in HeLa cells could be approximated by treatment with 5 mM histidine and 0.5 mM tryptophan in modified HEPES buffer (pH 7.4). The detection conditions were similar to those in live-cell fluorescence measurements. The mitochondrial tryptophan levels were estimated by fitting with purified protein titration curves at pH 8.0.

### Tryptophan Quantification with High Performance Liquid Chromatography

HPLC method was used to accurate quantify intracellular tryptophan levels in HeLa cells. We extracted the intracellular metabolites using 80% pre-cold methanol, evaporated the supernatant, and dissolved the samples in 100 µL of 90% acetonitrile to remove proteins before LC/MS quantification. The quantification process was carried out using a UHPLC system (1290 series; Agilent Technologies, USA) coupled with a quadruple time-of-flight mass spectrometer (TripleTOF 6600, AB SCIEX, USA). The obtained results were analyzed using the Agilent MassHunter Qualitative Analysis Navigator (B.08) software.

Mitochondria were extracted by a mitochondria isolation kit (Beyotime) and protein concentration was quantified with a BCA kit (Mxbioscience LLC), followed by LC/MS measurement. The protein amount was converted to mitochondria volume as reported ^7^.

The plasma tryptophan level was quantified with pre-column derivatization and reverse phase ultra-performance liquid chromatography (RP-HPLC)^34^. In brief, protein was precipitated by acetonitrile treatment in serum samples, and centrifuged at 13,000 rpm for 20 min. Derivatization of the supernatant was carried out at 55 °C for 10 min, followed by separation and quantitative analysis by external standard method with RP-HPLC.

### Statistical Analysis

Data are presented as the mean ± s.e.m. from at least three experiments. All statistical analyses and graphical representations were performed using R and Python. The figures were compiled and finalized using Adobe Illustrator 2020.

The normality of the data was tested using the Kolmogorov–Smirnov test. For comparisons between two groups, the two-tailed unpaired Student’s t-test was employed for data with a normal distribution in the majority of results, while the Mann-Whitney test was used for data with a non-normal distribution. The specific sample size (n numbers) and P values of each figure in the zebrafish and human serum sample experiments are listed in the corresponding figures or legends.

## Notes

### Competing Interest Statement

The authors have declared no competing interest.

