## supplementary Figures and Tables for "Unveiling Tryptophan Dynamics and Functions Across Model Organisms via Quantitative Imaging"

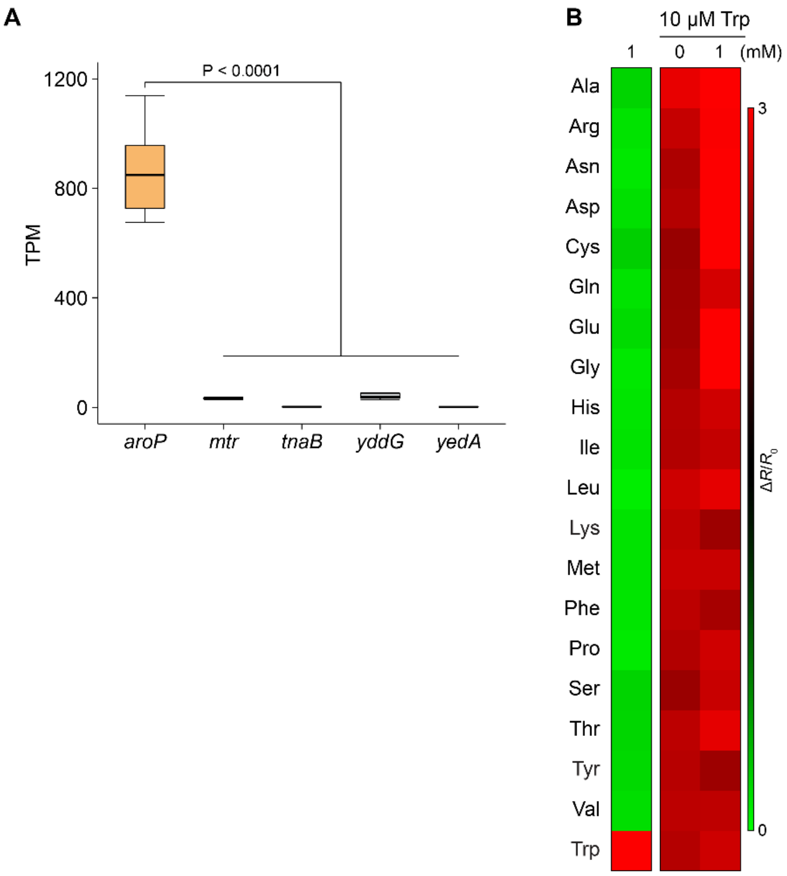


**Figure S1, related to Figure 1. Detection of tryptophan uptake in living bacteria. (A)** The expression level of tryptophan transporters on bacteria plasma membrane (n = 15). Data were taken from previous reports[1]. **(B)** Living bacteria expressing GRIT sensor responded to 1 mM different amino acids in the absence or presence of 10 μM Trp. n = 3 - 5 experiments. Data shown as mean ± s.e.m. Two-tailed unpaired student's t test for A.

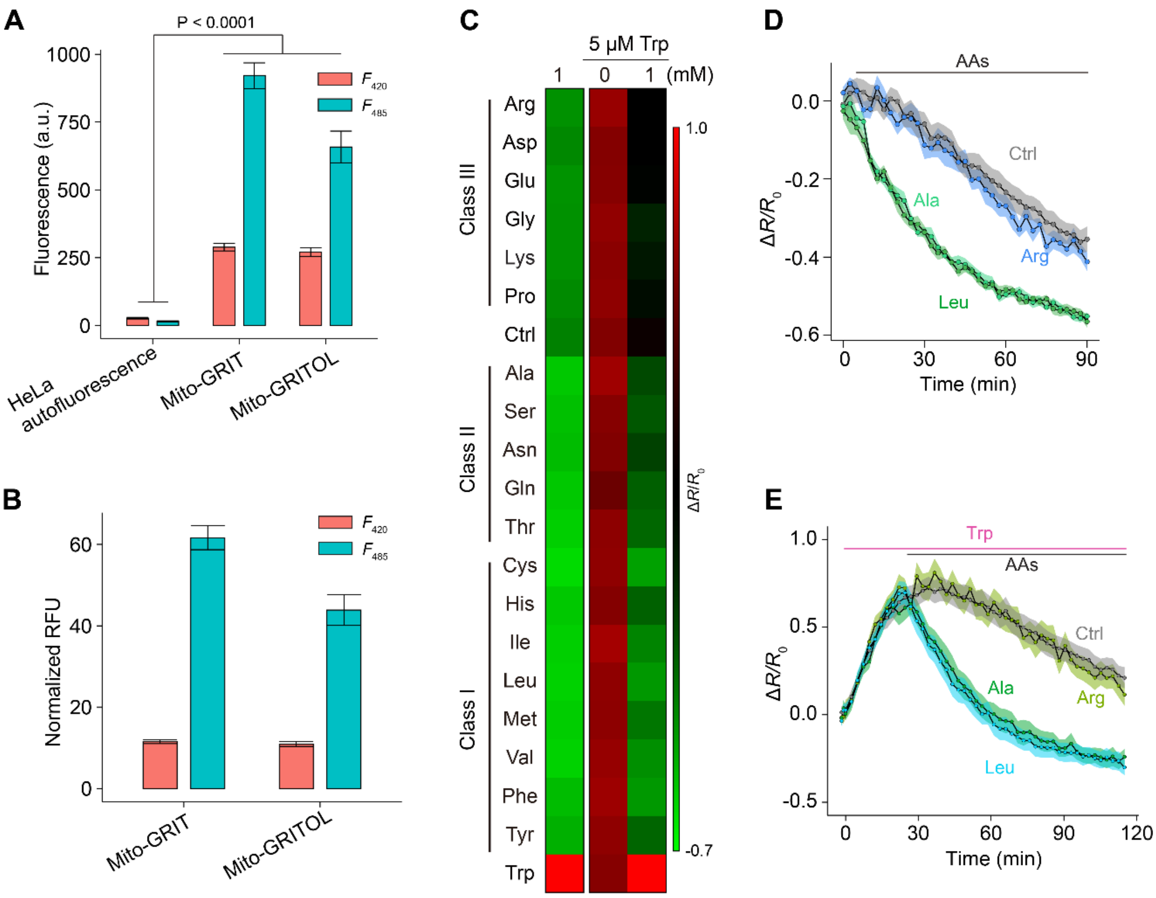


**Figure S2, related to Figure 2. Detection of tryptophan dynamics in mitochondria of HeLa cells. (A)** Raw dual excitation fluorescence intensities were measured in both HeLa cells and cells expressing either the mito-GRIT sensor or mito-GRITOL sensor. **(B)** The fluorescence values of mito-GRIT or mito-GRITOL positive HeLa cells were normalized to those of empty cells. The data presented in B are derived from A. **(C)** HeLa cell expressing mitochondrial GRIT responses to the indicated concentration of amino acids in the absence and presence of 5 μM Trp. **(D)** The responses of the mitochondrial GRIT sensor upon the addition of 1 mM indicated AAs. **(E)** Representative fluorescence traces of mitochondrial GRIT sensor following the addition of 5 μM Trp and the subsequent addition of 1 mM indicated amino acids.

Data shown as mean ± s.e.m. n = 3 - 5 independent experiments. Two-tailed unpaired student’s t test for A.

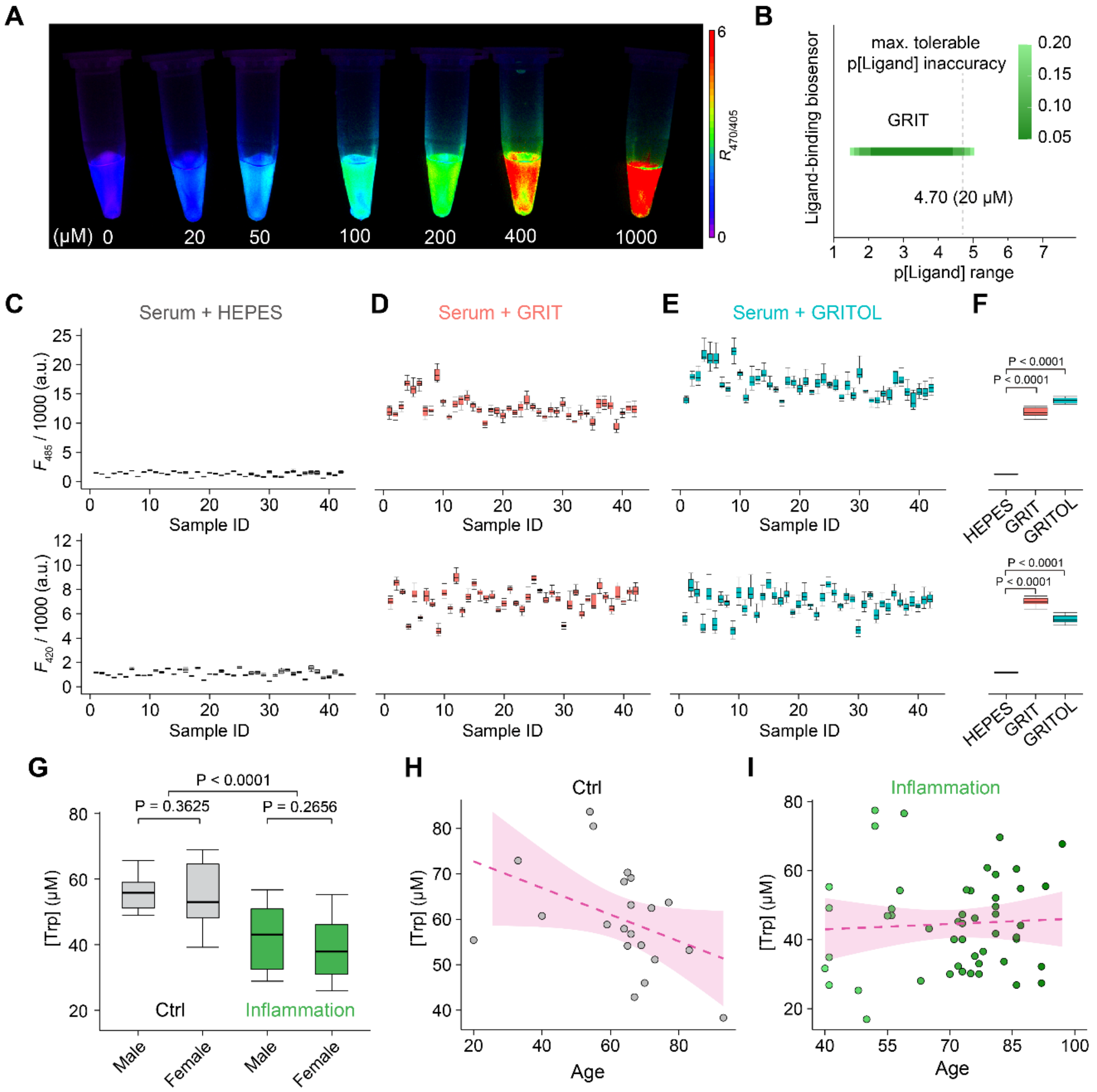


**Figure S3, related to Figure 3. Detection of tryptophan levels in the serum samples from inflamed individuals. (A)** Pseudocolored ratiometric images depict the response of 1 μM GRIT sensor to varying tryptophan concentrations. **(B)** The predicted measurement accuracy of the GRIT sensor across a wide range of tryptophan concentrations evaluated using the SensorOverlord application. **(C-E)** Original excitation fluorescence intensities ($F_{485}$, upper; $F_{420}$, down) of serum samples mixed with HEPES buffer (C), GRIT sensor (D), and GRITOL sensor (E). **(F)** The comparison of mean excitation fluorescence intensities ($F_{485}$, up; $F_{420}$, down) of serum samples with HEPES buffer, GRIT sensor, and GRITOL sensor. **(G)** Tryptophan concentrations of human serum samples in the control group (grey) and inflammation group (green) based on gender (G). Subgroup sizes range from n = 7 to 36. **(H-I)** Relationships between tryptophan concentrations in human serum samples and ages of control group (H) or inflammation group (I). The dashed magenta line represents a linear fit of the data, with the shaded area indicating the fitting confidence interval.

Data shown as mean ± s.e.m. Two-tailed unpaired student's t test for F-G.

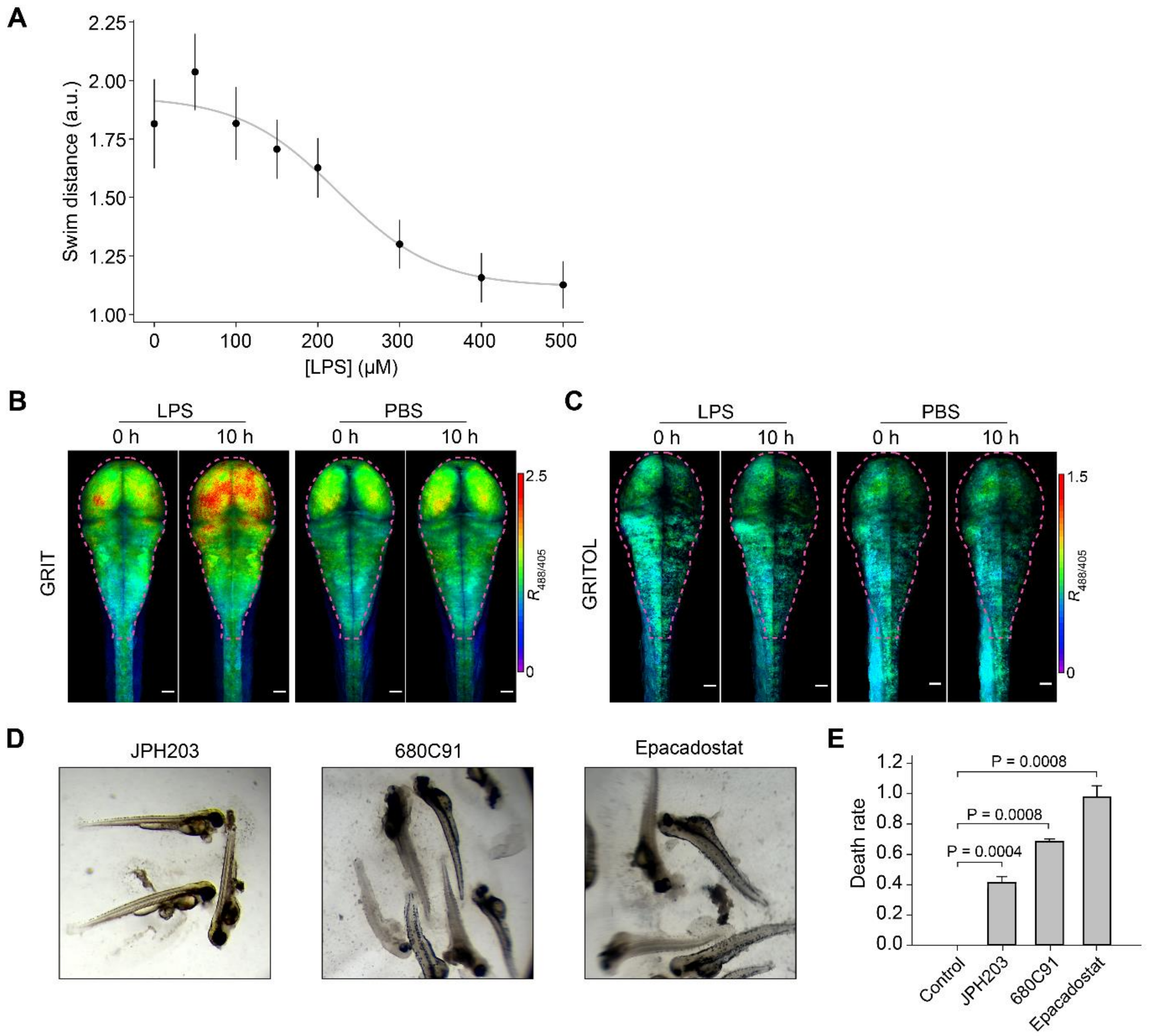


**Figure S4, related to Figure 4. Zebrafish sleep changes during inflammation, detection of metabolites levels in zebrafish larvae, and the death rate post tryptophan metabolism related inhibitors treatment. (A)** The dose-response curve of mean swim distance at night of zebrafish treated with different concentrations of LPS. **(B-C)** Representative images (maximum projection along the z-axis) of GRIT (A) or GRITOL (B) signals in the brain (dashed line in magenta) of zebrafish treated with LPS or PBS. Scale bars, 50 μm. **(D-E)** Representative images (D) and death rates (E) of larval zebrafish treated with 50 μM LAT1 inhibitor JPH203, TDO inhibitor 680C91, IDO inhibitor Epacadostat for 10 hrs.

Data shown as mean ± s.e.m. n = 3 - 5 independent experiments. Mann-Whitney U test for E.

**Table S1** | Quantifications of tryptophan level in bacteria, mammalian cells, zebrafish, and human serum. [a]Bacterial tryptophan level calibrated in HEPES buffer in the absence (upper) or presence (down) of tryptophan; [b]Reported tryptophan level in bacteria, taken from[2]; [c]Reported tryptophan level and uptake rate in cultured HeLa cells, taken from[3]; [d]Reported tryptophan level in cultured COS-7 cells, taken from[4]; [e]Mitochondrial tryptophan levels in HeLa cells measured in HEPES buffer; [f]Reported mitochondrial tryptophan level in cultured HeLa cells, taken from[5]; [g]Mean tryptophan levels of 42 identical serum samples measured by GRIT or HPLC/MS; [h]Mean tryptophan levels of serum samples from control patients (25) and inflamed patients (52) measured HPLC/MS; [i]Reported tryptophan level in human serum, taken from[6];

| | [Trp] (μM) | | | Uptake rate |
|---|---|---|---|---|
| | GRIT | HPLC | Reference | (μM/min) |
| Bacteria | 10.6 ± 3.7[a]<br>192 ± 23.9[a] | ND | 12[b] | <br>16.5 ± 1.4 |
| HeLa cytosol | 157.2 ± 17.4[c] | 207.3 ± 8.2[c] | 270 – 600[d] | 52.6 ± 13[c] |
| HeLa mitochondria | 112.4 ± 10.0[e] | 108.0 ± 10.3[e] | 29 – 77[f] | 9.6 ± 1.2 |
| Human serum | 51.9 ± 1.8[g] | 52.3 ± 2.0[g]<br>44.7 ± 4.7[h]<br>59.6 ± 3.6[h] | 54.5 ± 9.7[i] | ND |

**Table S2** | The effect of 19 amino acids on the tryptophan dynamics in mitochondria of HeLa cells. The rate is defined by tryptophan concentration decrease per minute (μM/min).

| Conditions | | Mitochondria |
|---|---|---|
| | Control | 0.80 ± 0.13 |
| Class I | Cys | 1.13 ± 0.07 |
| | His | 1.03 ± 0.04 |
| | Ile | 1.09 ± 0.05 |
| | Leu | 1.09 ± 0.07 |
| | Met | 1.05 ± 0.07 |
| | Phe | 1.01 ± 0.08 |
| | Tyr | 0.92 ± 0.07 |
| | Val | 1.09 ± 0.05 |
| Class II | Ala | 1.02 ± 0.05 |
| | Ser | 1.03 ± 0.08 |
| | Asn | 1.00 ± 0.05 |
| | Gln | 1.02 ± 0.05 |
| | Thr | 1.07 ± 0.08 |
| Class III | Arg | 0.84 ± 0.13 |
| | Asp | 0.79 ± 0.11 |
| | Glu | 0.81 ± 0.11 |
| | Gly | 0.75 ± 0.08 |
| | Lys | 0.71 ± 0.07 |
| | Pro | 0.72 ± 0.08 |
